# High-frequency extracellular spiking in electrically-active cancer cells

**DOI:** 10.1101/2024.03.16.585162

**Authors:** Rustamzhon Melikov, Francesco De Angelis, Rosalia Moreddu

## Abstract

Microelectrode matrices have been extensively used in the past 30 years as a reliable platform to record the extracellular activity of cells traditionally regarded to as excitable, i.e. brain cells and muscle cells. Meanwhile, fundamental biology studies on cancer cells since the 1970s reveal that they exhibit altered functionalities of ion channels, membrane potentials, metabolism, and communication mechanisms when compared to their healthy counterparts. In this work, we present for the first time the presence of extracellular voltage spikes occurring at high frequencies (0.1-3.5 kHz) in breast cancer cells, resembling the ones observed in excitable cells, and the possibility to record them with 30 µm TiN microelectrode matrices. These preliminary findings may open a new path for exploration in a variety of research fields, enabling the access to bioelectrical dynamics in cancer cells and cell networks, targeting bioelectricity as a tool in anticancer drug development and cancer diagnostics, as well as provide a market expansion for companies commercializing microelectrodes and microelectrode recording systems.

## Introduction

Bioelectricity is known to play a key role in regulating and driving crucial life processes at all scales: from subcellular structures, to cells and cell networks, tissues, and organs.^1^ The field of electrophysiological recording has evolved substantially over time in all these directions, translating these electric charges into actionable information in healthcare and drug testing.^2-5^ In the last 20 years, a myriad of technologies have been developed to access electrical information from cells using a variety of materials and techniques, each with its advantages and drawbacks. These include purely electrical methods, such as patch-clamp technique, microelectrodes,^6^ 3D nanoelectrodes,^7^ impedance spectroscopy,^8^ as well as optical probes, such as fluorescent dyes^9^ and genetically-encoded voltage indicators.^, 10^ Recently, combined approaches have been demonstrated. Among them, the use of electrochromic materials^11^ and charge-mirroring nanoelectrodes.^12, 13^ However, nearly all of the studies present in the existing literature have been focused on recording action potentials and extracellular signals in excitable cells, i.e. brain cells and muscle cells.^13-15^

Despite the utility of all these advancements, this research trend in electrophysiological recording technologies disregards the crucial role that electric charges play in all the remaining types of cells, i.e. the vast majority of biological cells existing in nature. Bioelectricity is known to be involved in regulating homeostasis, cellular migration,^5, 16, 17^ communication,^18^ proliferation,^19^ mutation,^20^ genetic expression,^21^ disease insurgence,^22^ DNA methylation^23^ and cell cycle progression, mythosis,^24^ and more. Electric gradients, resulting in potential differences at different levels (e.g., subcellular,^25^ cellular,^26^ and transepithelial^5^) can be generated by multiple biophysical processes, including passage of ions in and out of the cellular membrane,^27^ network currents given by cellular communication (e.g., gap junctions,^28^ paracrine signaling^29^), and surface charges^30^ resulting from altered cell metabolism.^1^

Cancer cells exhibit altered metabolism and communication compared to their healthy counterparts,^29, 31-33^ and this preliminary biological knowledge has driven the first investigations in the field of cancer bioelectricity. Most in vitro studies seem to be looking at comparing static membrane potentials of different cancer types or subtypes, or regard the biological studies of ion channels expression.^34^

Only three recent studies hint at the possibility of performing cellular electrical recordings in cancer cells.^4, 35-37^ However, both record signals occurring at low frequencies. M. Ribeiro *et al*. present large-area, low-impedance microelectrodes to record the electrical activity of breast cancer cell populations, registering spikes with duration above 400 ms.^35^ These signals were interpreted as network signals. The same approach was previously adopted in prostate cancer cells.^4^ The other main work in the field of cancer electrical recording was reported by P. Quicke *et al*.,^37^ where dynamic voltage imaging was conducted in cancer cells using bespoke genetically-encoded voltage indicators. However, the disadvantages of voltage imaging lie on i) the altered durations of the recorded signals, which means that if the duration is unknown, as it is the case for all cells other than neurons and cardiomyocytes, the information collected is only partial, and ii) the usual risk of altering cellular behavior by direct exposure to binding molecules. Signals recorded by voltage imaging had comparable durations to the ones recorded in cell populations.

Nevertheless, both works convey a precious information: the electrical activity of these cells is visibly dynamic. Following this valuable insight, we performed single (or few) cells electrical recording of breast cancer cells using 30 µM TiN microelectrode matrices (60 electrodes), with data acquisition both at low (1-100 Hz) and high (100-3500 Hz) frequencies. Interestingly, we found out that i) breast cancer cells not only exhibit cohort signaling, but also single-cell signals recordable with tiny electrodes just as typically done in excitable cells; ii) breast cancer cells (and, we expect, many other cancer cell types), exhibit not only low-frequency signals, as demonstrated previously in the reviewed works, but also and most importantly high frequency spiking and bursting that clearly resemble the ones observed in excitable cells.

## Results

We characterized four subtypes of triple negative breast cancer cells of two different families: MDA-MB-231 and MDA-MB-468, HCC-1937 and HCC-1143. MDA-MB-231 and HCC-1937 are known to possess higher metastatic ability compared to MDA-MB-468 and HCC-1143, respectively. All cells exhibited visible activity both at low frequencies (1-100 Hz) and high frequencies (100-3500 Hz).

MDA-MB-231 cell line (**Figure 1**) exhibited the highest activity and remarkable network synchronicity as visible in the Raster plot in **Figure 1A**. Average spike durations of 8-10 ms and 500-800 ms were calculated at high and low frequencies, respectively. Sample spikes are displayed in **Figure 1B. Figure 1C** display whole recordings (6 mins) of 3 selected channels. Channel 14 and 21 exhibited synchronicity with 19 and 16 more channels of the matrix, respectively. Amplitude of the recording was cut for visualization of spikes in the most occurring amplitude range. All channels recorded voltage spikes between 100 and 500 mV, with occasional higher amplitude spikes reaching up to few mV. Channels with an electrical activity of higher entity compared to all the other electrodes of the matrix were identified in each independent culture, leading to a speculation on the possible presence of a triggering or regulating role of one or few cells in the network. **Figure 1D** presents a sample case of whole-network synchronicity of a spiking event at high frequencies (45 channels) with amplitudes ranging from 200 µV to 1.5 mV and durations of 10 ms. Hybrid signals resembling 3D biological constructs have been identified, together with clean individual spikes. MDA-MB-231 cell line also exhibited bursting activity, with spikes in bursts of duration comparable to that observed in individual spikes (10 ms), but lower amplitude. Different types of bursts were identified: i) bursts containing clean consecutive spikes (**Figure 1E**); ii) bursts containing a continuous hybrid signals or a sequence of hybrid events (**Figure 1F**). Similarly, observed spikes were either individual, as shown earlier in Figure 1B, or hybrid as in Figure 1F.

**Figure 1.**
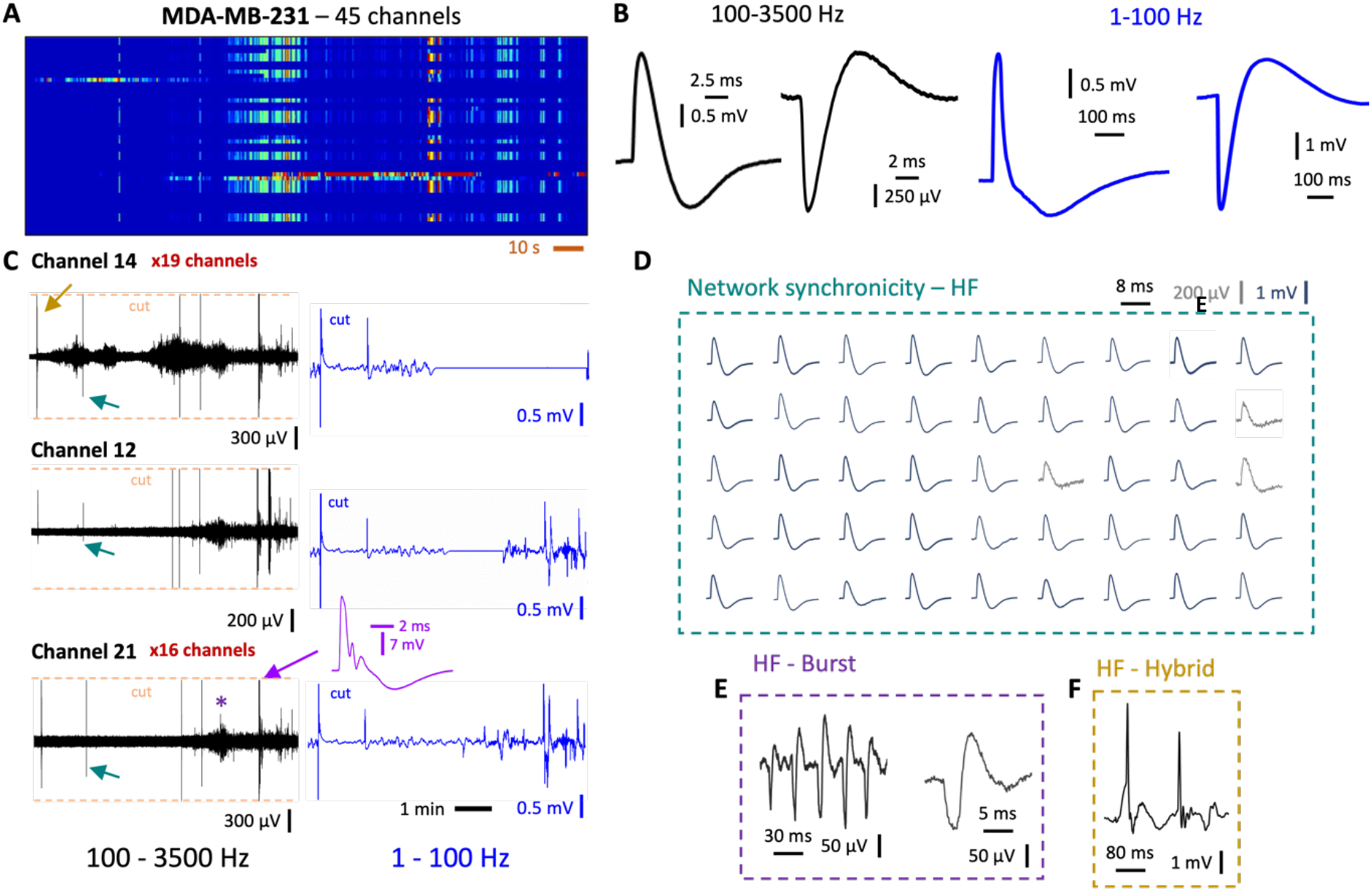
Electrical activity of the MDA-MB-231 cancer cell line. A) Raster plot of 45/60 firing channels (3 min). B) Sample high frequency and low frequency spikes observed in MDA-MB-231 cell line. C) 6 minute recordings of 3 selected channels at high and low frequency (simultaneous). D) Network synchronicity in 45 channels. E) Busting activity characterized by individual, clean consecutive spikes as extracted from purple star (channel 21). F) Sample of hybrid spiking events happening in channel 14 in the point indicated by the golden arrow.

Figure 2 displays the results obtained with MDA-MB-468 cells, possessing a lower metastatic ability compared to MDA-MB-231. The sample Raster plot in **Figure 2A** shows a remarkably lower activity with respect to the MDA-MB-231 cell line, in terms of active channels and spike amplitude, as well as a visibly lower synchronicity. Spike duration was comparable to the MDA-MB-231 cell line both at high and low frequencies (**Figure 2B**). **Figure 2C** presents 3 selected channels of the network. A bioelectrical hallmark of this cell line was found to be the presence of voltage microbursts (50-80 µV) with shape as depicted in **Figure 2D**, total duration of 5 ms. Corresponding bursts were identified at low frequencies (**Figure 2E**) with falling peaks of 50 ms duration.

**Figure 2.**
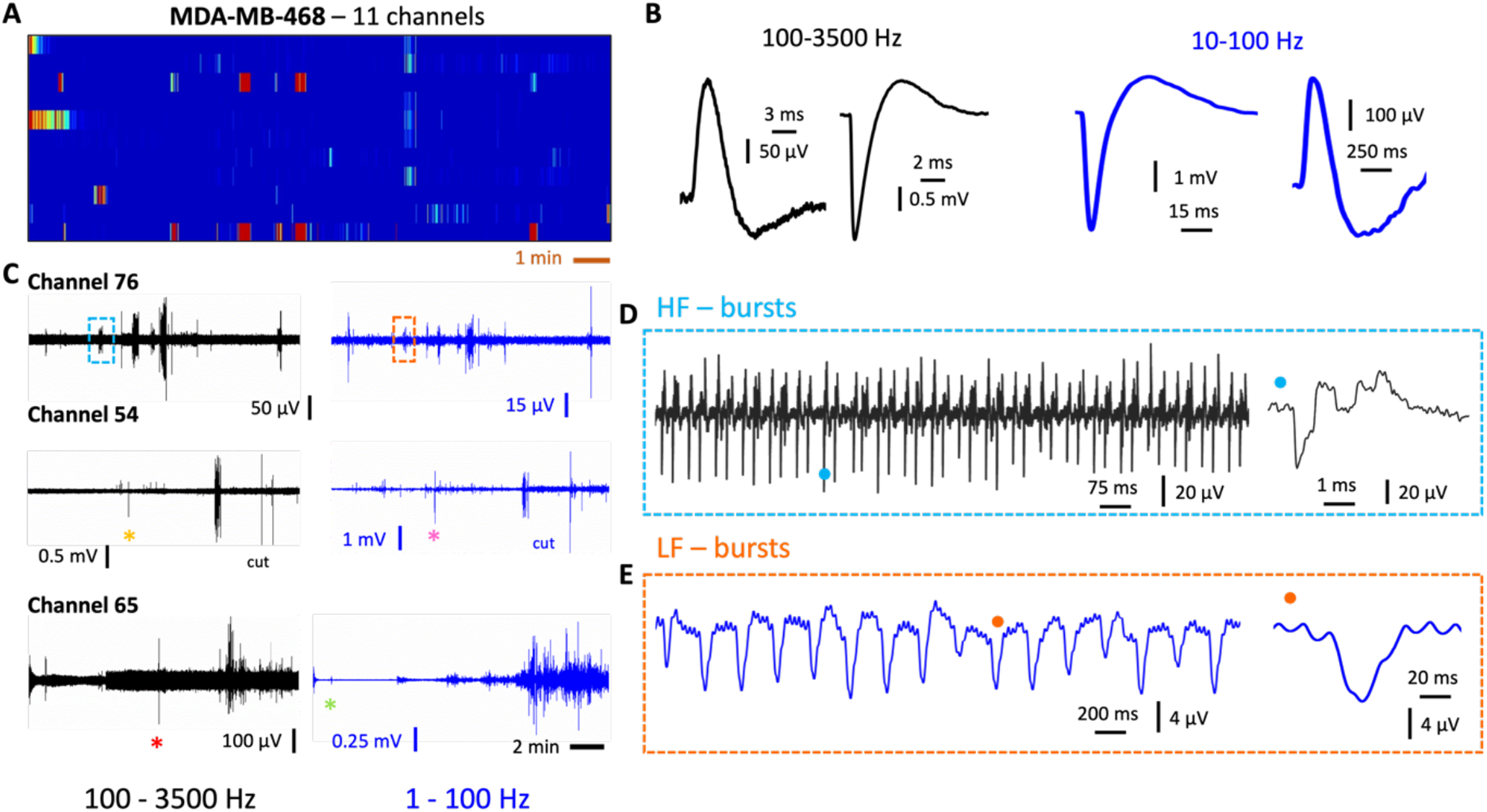
Electrical activity of the MDA-MB-468 cell line. A) Raster plot of 11/6**I** 0 firing channels. B) Sample high frequency and low frequency spikes observed in MDA-MB-468 cell line. C) 10 minute recordings of 3 selected channels at high and low frequency (simultaneous). D) Microbursts extracted from the light-blue dashed square area in Channel 76. E) Bursting at low frequency extracted from orange star in channel 54.

Two cell lines of the HCC family were characterized, differing for their metastatic ability. **Figure 3** shows sample recordings obtained with the HCC-1937 cell line, with high metastatic potential, reporting 11 selected channels of the network. Here, a lower number of active channels (hence, cells) were identified, compared to the MDA-MB-231 highly metastatic cell line. However, a higher spike rate per channel was calculated. The Raster plot in **Figure 3A** shows one highly active, bursting cell, and a noticeable synchronicity with varying amplitude in different channels. This difference compared to what observed in MDA-MB-231 may suggest the presence of coordinated signals among different cells of the network (HCC-1937) Vs a whole network activity (MDA-MB-231). High frequency spikes with duration 10-12 ms were recorded consistently; low frequency activity comprised peaks with variable duration, from 50 ms to 700 ms (**Figure 3B**). **Figure 3C** displays 5 selected channels exhibiting both spiking and bursting activity, with spike amplitudes ranging from 200 µV to 5 mV. Bursts were mainly composed of hybrid signals with occasional individual peaks (**Figure D**). The pink square in Figure **Figure 3E** comprehends spikes extracted from channel 43 in **Figure 3C**, showing a consistent duration of approximately 10 ms. Bursts were also identified at low frequencies with durations of 300 ms, distinguishable from noise, as displayed in **Figure 3F. Figure 3G** presents individual low-frequency peaks extracted from channel 58, with total durations of 600 ms.

**Figure 3.**
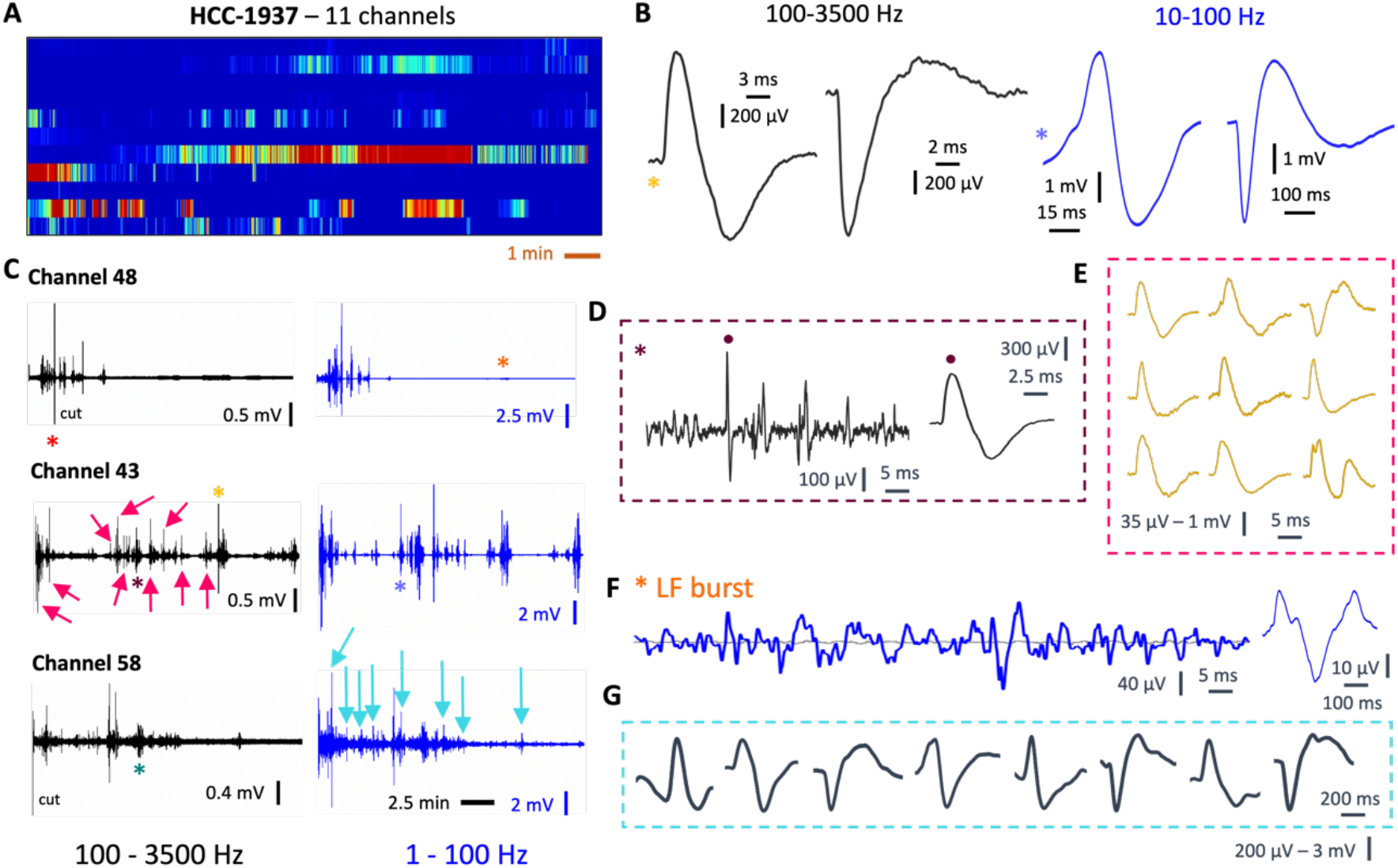
Electrical activity of the HCC-1937 cell line. A) Raster plot of 11/60 firing channels. B) Sample high frequency and low frequency spikes, observed in HCC-1937 cell line. C) 10 minutes recordings of 3 selected channels at high and low frequency (simultaneous). D) Hybrid bursting activity extracted from channel 43 (purple star), similar to what observed in the bursting areas of other channels. The inset shows a clean extracted spike. E) High frequency spikes as extracted from highly active channel 43 and indicated by fucsia arrows in figure. F) Low-frequency bursts distinguishable from baseline noise, extracted from channel 48 (orange star). G) Low frequency spikes as extracted from channel 58 and indicated by light-blue arrows.

Figure 4 presents the characterization of the HCC-1143 cancer cell line, sharing the same family of the HCC-1937 line but lower metastatic ability. As observed in the MDA-MB family, a lower metastatic ability correlated to a lower electrical activity. However, HCC-1143 line displayed visible synchronicity and a higher spike-to-bursts ratio compared to the MDA-MB-468 cell line of the previous family, as visible in the Raster plot presented in **Figure 4A. Figure 4B** shows high and low frequency spikes of durations 10-12 ms and 120-600 ms, respectively. **Figure 4C** presents whole recordings (10 min) from 8 channels, displaying high synchronicity both at low and high frequencies.

**Figure 4.**
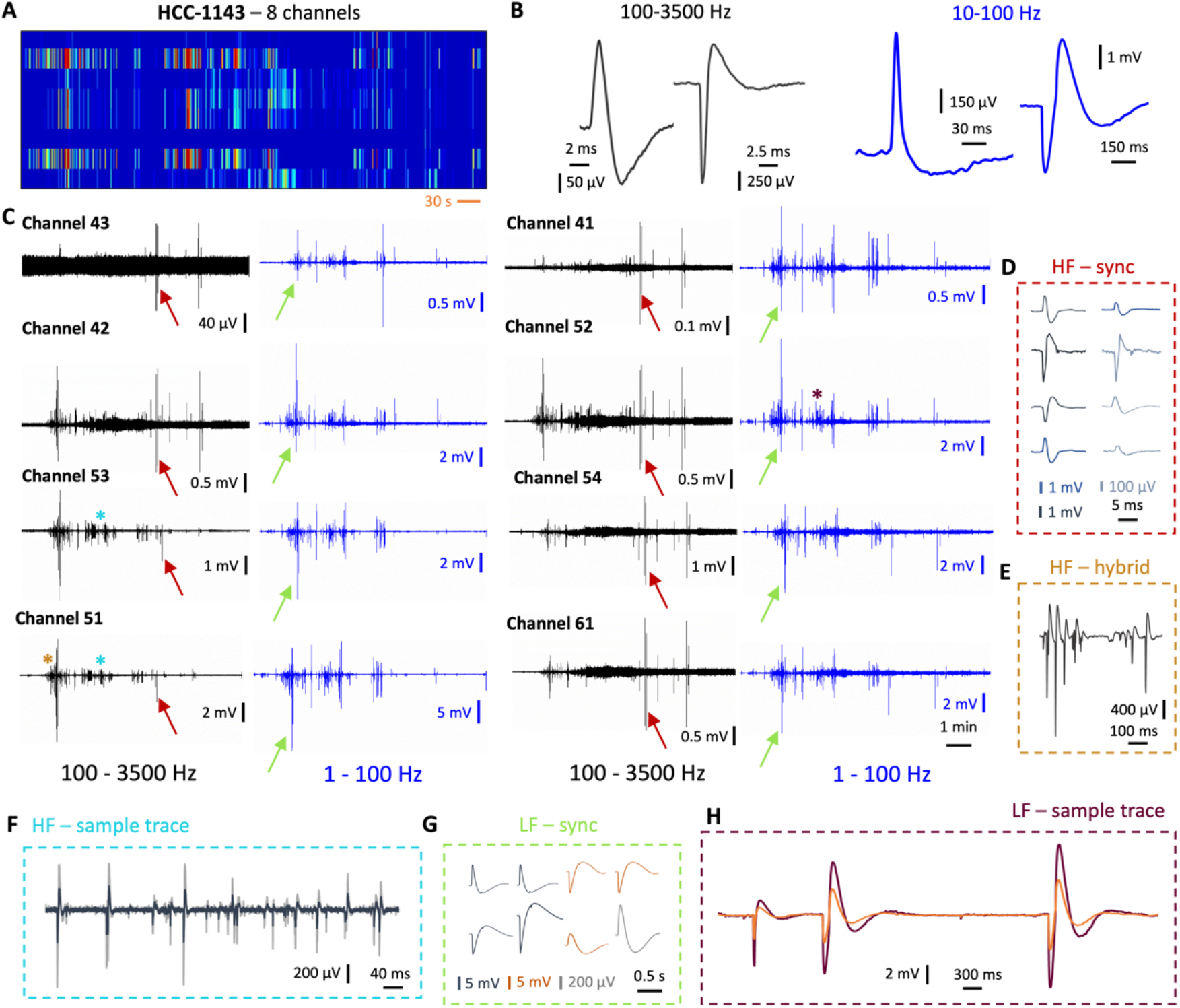
Electrical activity of the HCC-1937 cell line. A) Raster plot of 8/60 firing channels. B) Sample high frequency and low frequency spikes, rising and falling, observed in HCC-1143 cell line. C) 10 minute recordings of 8 selected channels at high and low frequency (simultaneous). D) High-frequency synchronic spiking in 8 channels, as indicated by the red arrows. E) Hybrid consecutive bursts extracted by channel 51 (golden star). F) High frequency synchronic trace extracted from channels 51 and 53 (light blue stars). G) Low-frequency synchronic spiking in 8 channels, as indicated by the green arrows. H) Low-frequency sample trace, extracted from channel 52 (purple star).

Sample synchronous high frequency spikes are presented in **Figure 4D**, with amplitudes ranging from 100 µV to 5 mV and durations of 8 ms. The gold square in **Figure 4E** reports a hybrid bursting activity extracted from channel 51, with amplitudes up to 3 mV. High frequency sample traces (**Figure 4F**) report a snapshot on the activity of two selected channels. Synchronicity was observed also at high frequency, and extracted sample peaks of duration 1s are presented in **Figure 4G**. A low-frequency synchronic sample trace of two selected channels is also reported in **Figure 4H**.

Figure 5 presents a statistical comparison plot between the examined cell lines. Burst detection (**Figure 5A**) and spike detection (**Figure 5B**) analyses revealed higher counts in MDA-MB-231 and HCC-1937, the cell lines with the highest metastatic level. MDA-MB-231 cells were found to be the most active, firing an average of 60 spikes per second at the network level, followed by over 50 spikes/sec in HCC-1937, 25 spikes/sec in MDA-MB-231, and 8 spikes/sec in HCC-1143. Bursting activity followed the same trend, with the highest percentage of spikes in bursts found in MDA-MB-468 cells. The mean spike rate in bursts was found to be comparable in MDA-MB-231, MDA-MB-468, and HCC-1937 cells, and lower in HCC-1143 cells. Despite MDA-MB-468 exhibiting slightly higher electrical activity compared to HCC-1143, the number of active channels was found to be higher in HCC-1143 networks. Moreover, HCC-1937 was found to be the cell line with the highest spike-to-burst ratio.

**Figure 5.**
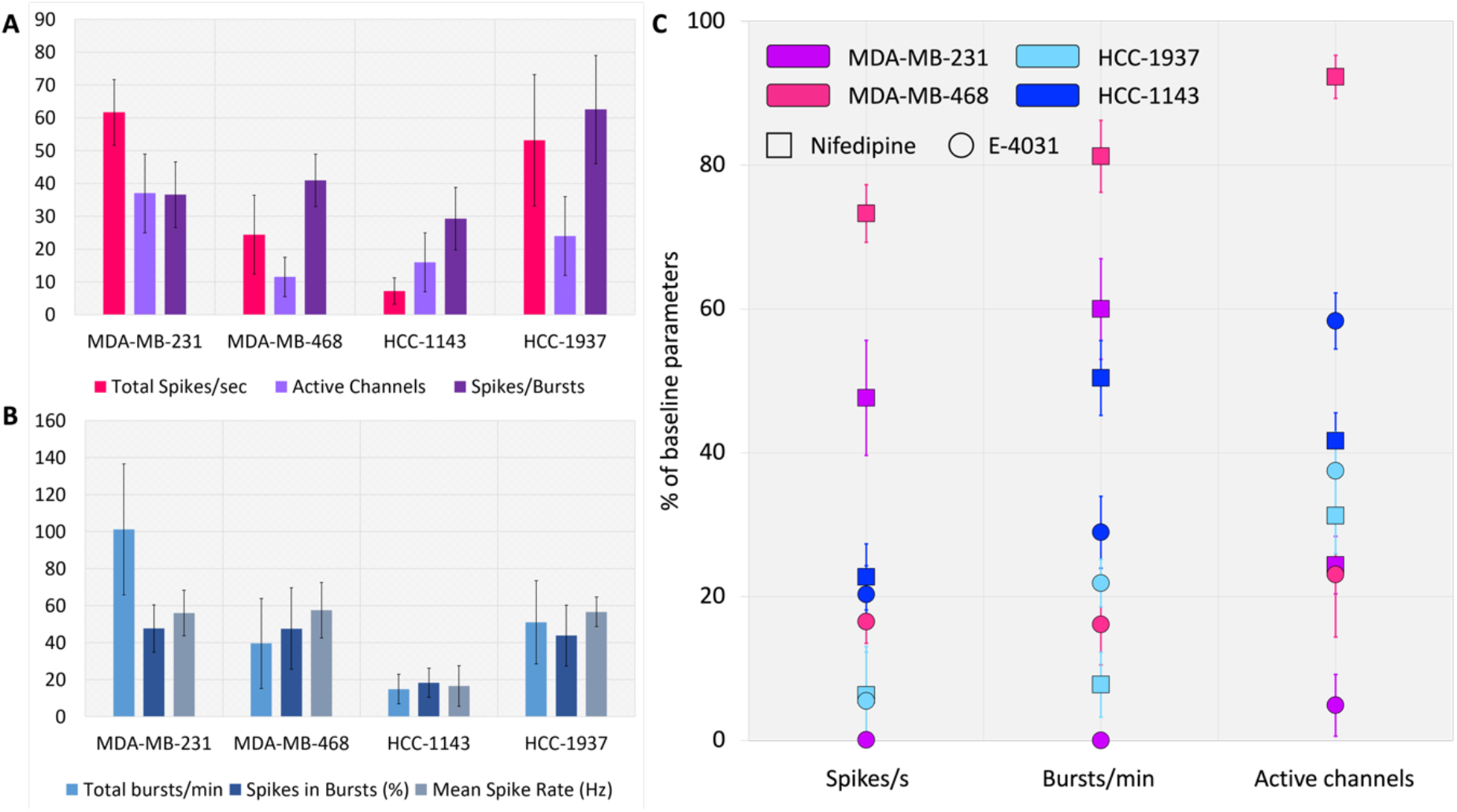
Comparison on the electrical activity observed in the 4 examined cancer cell lines and effect of selective channel blockage on spiking, bursting and network activity. A) Evaluation of the spiking activity of the four lines, with regards to spiking rate, number of active channels and spiking activity over bursting activity prevalence. B) Evaluation of the bursting activity of the four lines, with regards to bursting rate, percentage of spikes in bursts, and mean spike rate in bursts. C) Summary of selective h-ERG K^+^ and L-type Ca^2+^ channels blockage via E-4031 20µM and Nifedipine 20µM administration to the four cell lines on microelectrodes.

To confirm the biological nature of the recorded electrical activity and to evaluate the ion channel expression of the examined breast cancer cell lines, we selectively blocked h-ERG K^+^ channels and voltage-dependent L-type Ca^+^ channels by administering E-4031 and nifedipine, respectively. **Figure 5C** present a summary of the results. All cell lines exhibited a drop in electrical activity (spiking, bursting and active channels) following drug administration, with remarkably different levels of response. The graph in figure presents the percentage of spikes/s, bursts/min and active channels after drug administration, compared to baseline activity. Notably, all cells showed a more marked response to E-4031 with a reduction in spiking to 20% and below, and bursting to 30% and below. Nifedipine was the least effective in MDA-MB-468, where both spiking and bursting rate were recorded to be between the 70 and 80% to the baseline, followed by MDA-MB231 with a fall to the 50-65%. Nifedipine administration to cell line HCC-1143 resulted in a spiking fall to the 20% compared to baseline, and a bursting fall to the 50%. On the other hand, nifedipine nearly killed activity in highly metastatic HCC-1937 cells.

Sample Raster plots and sample recordings in baseline and following E-4031 or nifedipine administration are presented in **Figure 6. Figure 6A** reports sample results obtained in the cell line MDA-MB-231. Raster plots (grey) display a visible fall in both spiking and bursting after nifedipine administration, and null activity following E-4031 administration. **Figure 6B** reports results obtained with HCC-1143 cell line in the same conditions, resulting in a lower spike rate following nifedipine administration as well as E-4031 administration. None of the compounds killed electrical activity, suggesting a speculation on possible compensatory mechanisms or the absence of the targeted ion channels in HCC-1143. **Figure 6C** presents sample results obtained with MDA-MB-468 cell line, where both drugs reduced the spiking rate, bursting rate and number of active channels. Despite the decreased firing rate and number of active channels, occasional spikes with higher amplitudes were recorded following drug administration, compared to baseline signals. Sample results obtained with cell line HCC-1937 are presented in **Figure 6D**. Here, the number of active channels was found to be poorly altered, in favour of a visibly decreased spike amplitude following nifedipine administration, and an increased amplitude following E-4031 administration. Both drugs induced a remarkably lower firing rate, as described in the previous paragraph.

**Figure 6.**
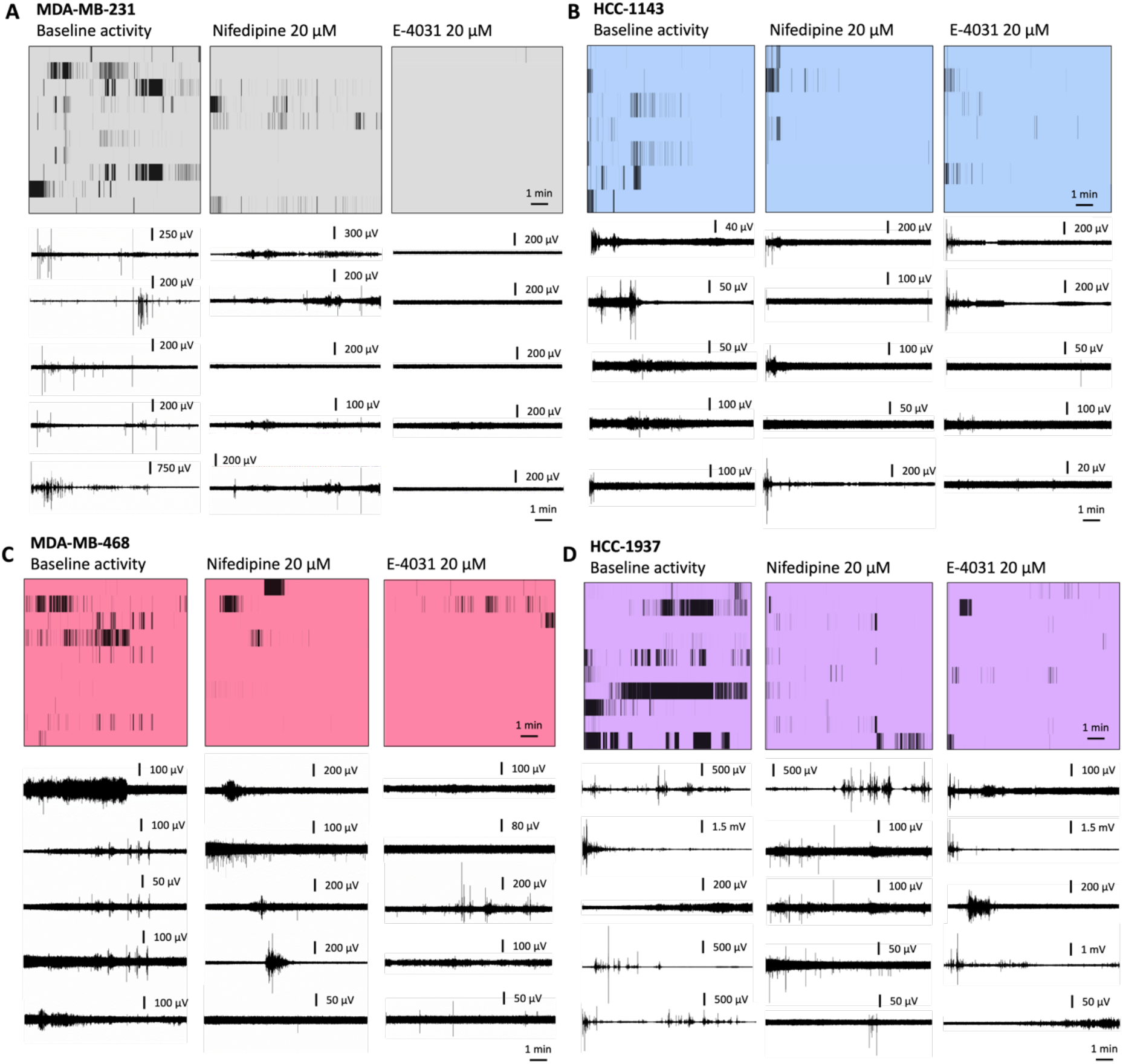
Sample results on the effect of selective channel blockage in the four examined cell lines: Raster plots and recordings of 5 selected channels of baseline activity, post-nifedipine (20µM) or post-E4031 (20µM) administration. A) MDA-MB-231; B) HCC-1143; C) MDA-MB-468; D) HCC-1937.

## Materials and Methods

### Cell culture

MDA-MB-231, MDA-MB-468, HCC-1143 and HCC-1937 cell lines were purchased from ATCC. All cells were expanded until passage 3 before seeding on chip, and all recordings were performed in cells at passages 4-12, to ensure reliability and reproducibility. Cells of the MDA-MB family were cultured in DMEM high glucose, 10% FBS, with L-glutamine, and 1% penicillin-streptomycin. Cells of the HCC family were cultured in RPMI 1640, 10% FBS, 1% penicillin-streptomycyn, L-glutamine.

### Cell seeding on chip

Electrodes were prepared equally for seeding all cell types. MEAs were sterilized for 30 minutes in a biohood, followed by drop casting of 0.01% poly-L-lysine solution (Sigma Aldrich) in the electrodes area. After one hour, chips were rinsed with sterile water and air-dryed. Cells were seeded at a concentration of 150-200k cells per chip to ensure confluency after 24-36 hours. Chips were stored in an incubator at 37°C and 5% CO_2_.

### Data acquisition

30 µm TiN 60 microelectrode arrays were purchased from Multi Channel Systems and seeded as described in the previous paragraph. The cell medium was changed four hours prior to each recording to ensure optimal viability. Electrical recordings were performed using a MEA-2100 system purchased from Multi Channel System, and data was acquired with the dedicated software at frequency ranges 1-100 Hz and 100-3500 Hz.

### Drug test

Channel blockers E-4031 and nifedipine were purchased from Sigma, aliquoted in DMSO at a concentration of 10 mM, and further utilized such to reach a concentration of 20µM in vitro for both compounds. Channel blockers were left acting for one hour, then rinsed with cell medium. Each experiment was performed in a separate MEA and statistical results were obtained by the data collected from5 independent measurements (two MEAs each), and 4 independent cell cultures per cell line.

### Data analysis

Data analysis was perfomed using the dedicated software Multi Channel Analyzer and MATLAB. Bursts were defined as events containing a minimum of 4 spiking events occurring with a maximum time interval of 300 ms. Threshold for rising-falling spike identification was defined automatically setting a standard deviation of 5%, then verified for each electrode to ensure a reliable calculation. Raster plots allowed to further visualize the spiking events.

## Conclusions

Our preliminary findings reveal that the examined breast cancer cells exhibit extracellular spikes occurring at high frequencies, that resemble the electrical activity observed in excitable cells. This data was acquired using commercial microelectrode arrays with a diameter of 30 µm to interface a few cells. The biological origin of the signals was further confirmed by pharmacological manipulation by selective K^+^ and Ca^2+^ channel blockage. By comparing the results obtained with different subtypes of cancer cells, we performed a preliminary analysis that correlates a higher fire rate to a higher metastatic ability. These results advance current knowledge limited to low-frequency signaling in cancerous cells, identify an accessible method to retrieve electrical information from cells beyond neurons and cardiac cells. As such, it may set the foundations to further developments in the fields of nanotechnology, drug testing, diagnostics, fundamental biology, and commercialization of microelectrodes.

## Acknowledgements

R.M. thanks the European Commission for a Marie Curie Individual fellowship (grant agreement number 101064443).

